# A Multi-Scale Modelling Framework of the Whole Human Body Based on Multi-Objective Functions

**DOI:** 10.1101/477703

**Authors:** Masood Khaksar Toroghi, William R Cluett, Radhakrishnan Mahadevan

## Abstract

Constraint-based multi-scale metabolic models are powerful tools to study complex biological systems like the human body, and to develop new treatment strategies for human diseases. They capture different scales of the system under study and provide the opportunity of understanding the interaction between multiple scales. In this paper, we have used our previously developed multi-scale whole body metabolism framework to establish a new approach to include multiple objectives of the human cells in computational analysis. The model has 3555 ordinary differential equations (ODEs) integrated with a genome-scale model of the hepatocyte, including 1826 biochemical reactions. The model has been solved for 74 objective functions simultaneously. Simulation results show that the integration algorithm has promise with respect to convergence, computational efficiency and response to perturbation. The results suggest that this method holds significant potential for the simulation of the range of metabolic phenotypes for mammalian cells where there are multiple objectives.

## Introduction

Constraint-based metabolic modelling approach has been widely used in the biomedical and life sciences research community [1]. These models have been used to characterize the metabolism of human cells, identify drug targets [2], study the pathophysiology of cancer cells [3], and identify new biomarkers in inborn errors of metabolism [4].

Research has shown that in the majority of metabolic models that are based on the constraint-based modelling approach, a single function of the living organism has been considered in the simulation analysis [5, 6]. In other words, in the analysis, a single objective function is solved at a time [7, 8]. However, biomedical research and experimental evidence have shown that mammalian cells are capable of performing multiple tasks at the same time. For example, the human liver cells have five tissue systems, which are the vascular system, hepatocytes and hepatic lobule, hepatic sinusoidal cells, biliary system, and stroma. In addition, there are many different cell populations in the liver, which are hepatocytes, endothelial cells, Kupffer cells and hepatic stellate cells [9]. As a result, the organ can perform several functions simultaneously, including synthesis of plasma proteins, detoxification of ammonia [10], drug metabolism [11], poisonous substance elimination, and glucose and ethanol metabolism [12,13] as depicted in Figure 1. Moreover, the liver also plays a significant role in an innate immune system that protects hosts from environmental threats, such as invading microorganisms, and physical and chemical stimuli [9]. In addition, the liver cells receive oxygen, nutrients and chemicals supplied by the circulation. Also, the liver processes blood by breaking down the nutrients, breaking down chemicals into solutes that can be used by other organs (e.g., the brain, heart, and kidney), storing incoming substances such as glucose, and regulating the level of most chemicals in the blood such as cholesterol and glucose [14,15,16]. This evidence indicates that including multiple functions of mammalian cells in our computational analysis is very important, because, first, it helps enhance the accuracy of prediction results, and second; it helps us to have a better understanding of metabolism.

**Figure 1:**
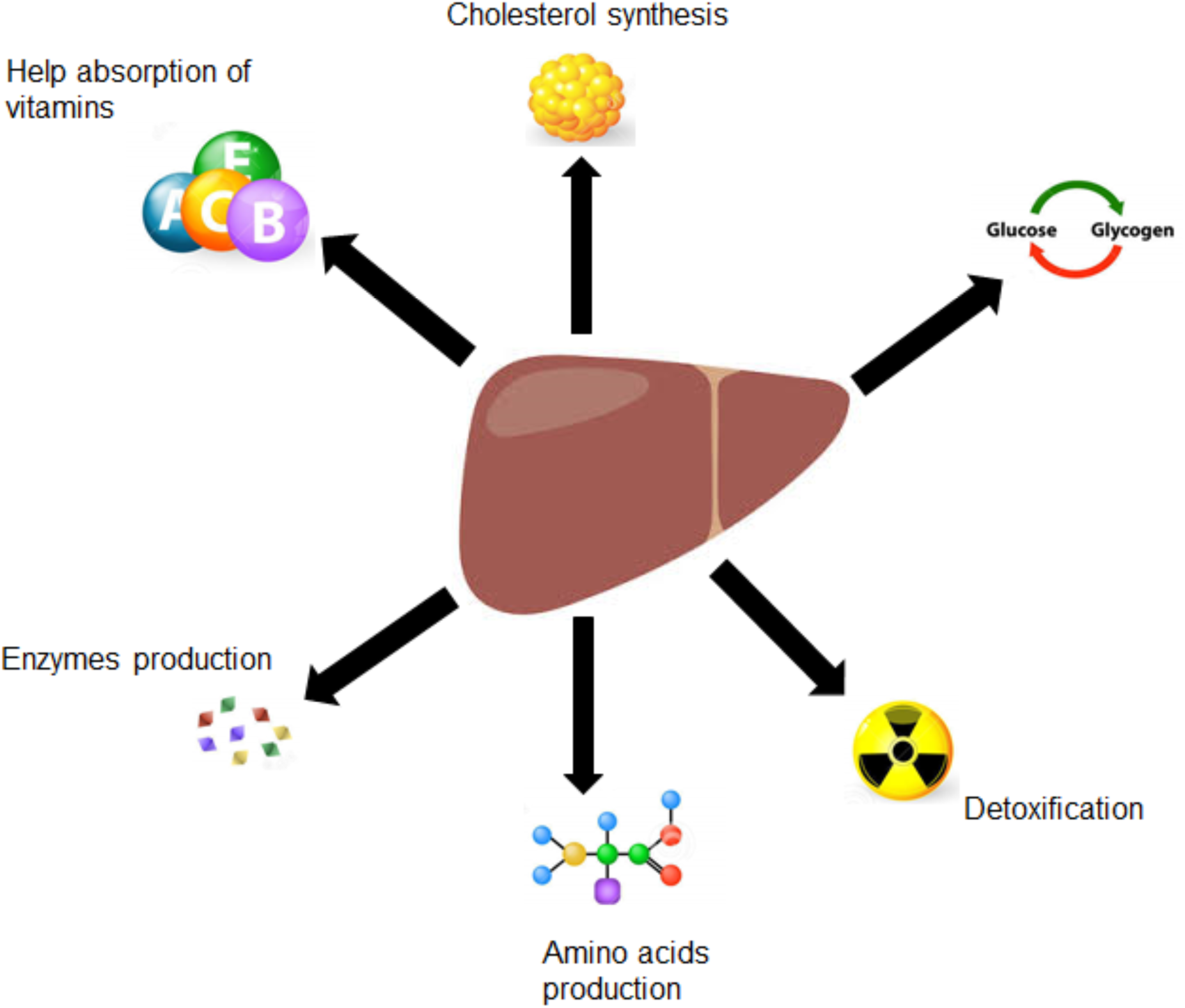
Liver functions. The liver plays a major role in metabolism with numerous functions in the human body, including detoxification of various metabolites, synthesis of protein, amino acid and cholesterol, help with the absorption of vitamins, and the production of enzymes.

In this paper, we have presented a new computational approach to include multiple functions of the human hepatocyte cells in the whole body metabolism framework. The algorithm is an extension of our previous study, presenting a multi-scale whole body metabolic model [5]. The computational efficiency of the integration algorithm has been demonstrated using several *in silico* studies.

## Results and Discussion

### 1. Model Simulation

To illustrate the computational efficiency and convergence of the multi-objective function framework, we have used the hepatocyte metabolic model developed by Gill et al. [12] as an example of human cells. In this network, cells can produce up to 74 metabolites. Here, we have considered the maximum number of objective functions (i.e., 74) for our analysis. To demonstrate the computational efficiency and convergence of the multi-objective framework, *in silico* analyses are performed for a limited simulation time. The model properties are presented in Tables A.1-A.3 in the Appendix.

Figure 2 demonstrates the concentration profiles for a number of consumed metabolites. As seen, metabolite concentrations converge to zero, because all the substrates are depleted after a period of time.

**Figure 2:**
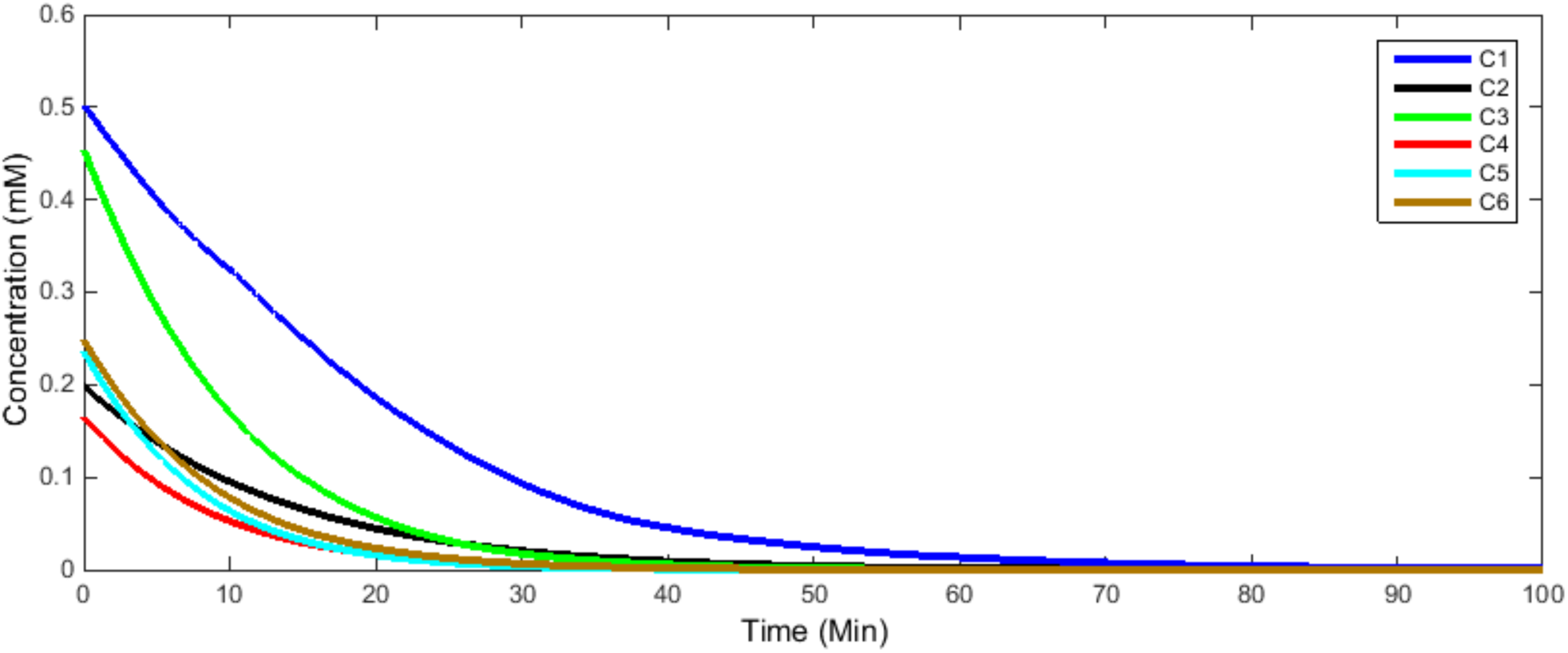
Concentration profiles for a number of consumed metabolites. Metabolite concentrations converge to zero, because all the substrates are depleted after a period of time.

In addition, Figure 3 shows the concentrations of the produced metabolites converge to new steady-state conditions.

**Figure 3:**
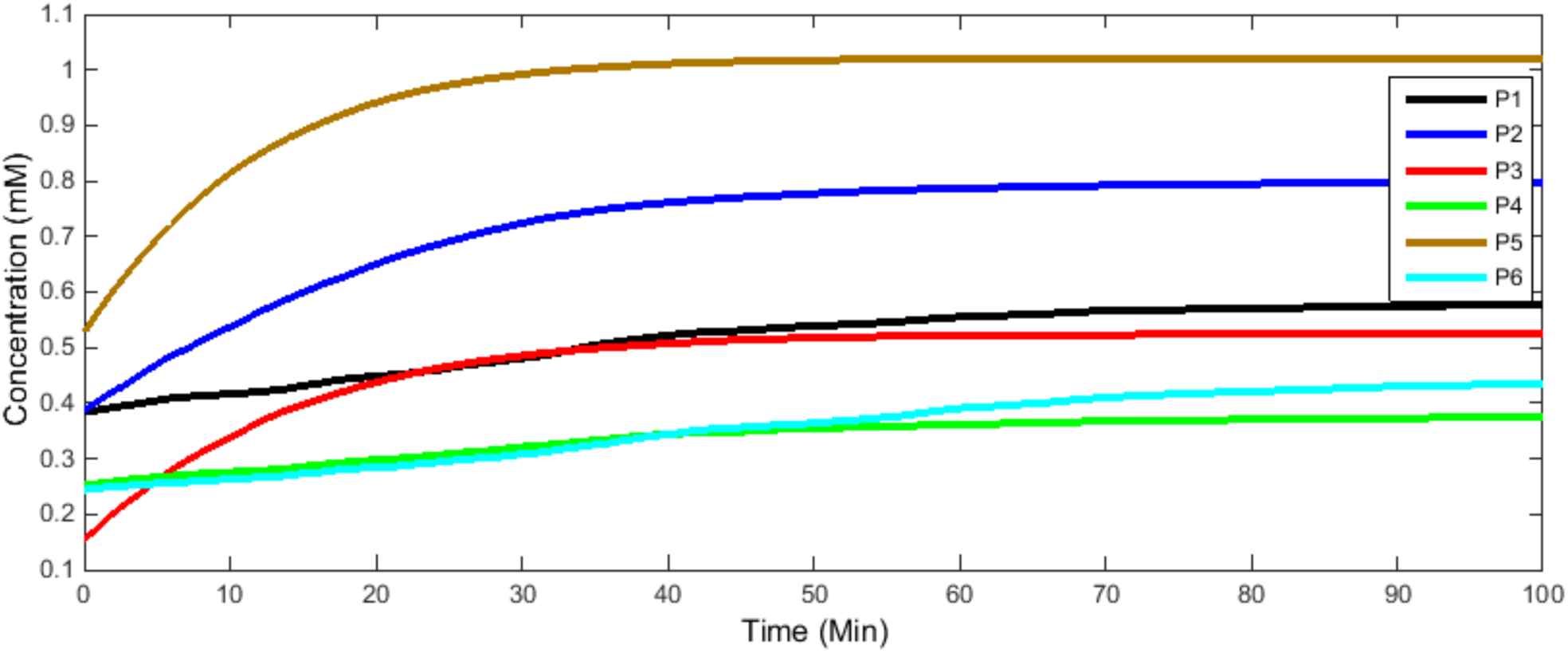
Concentration profiles for a number of produced metabolites. At steady-state conditions, the concentrations of the metabolites converge to new steady-state conditions.

The same analysis is conducted for metabolites that can be produced or consumed in the cell. As seen in Figure 4, the concentrations for the number of metabolites converge to zero (i.e., they are consumed by the cells), and concentration of two exchange metabolites reach new non-zero steady-state conditions (i.e., they are produced by the cells).

**Figure 4:**
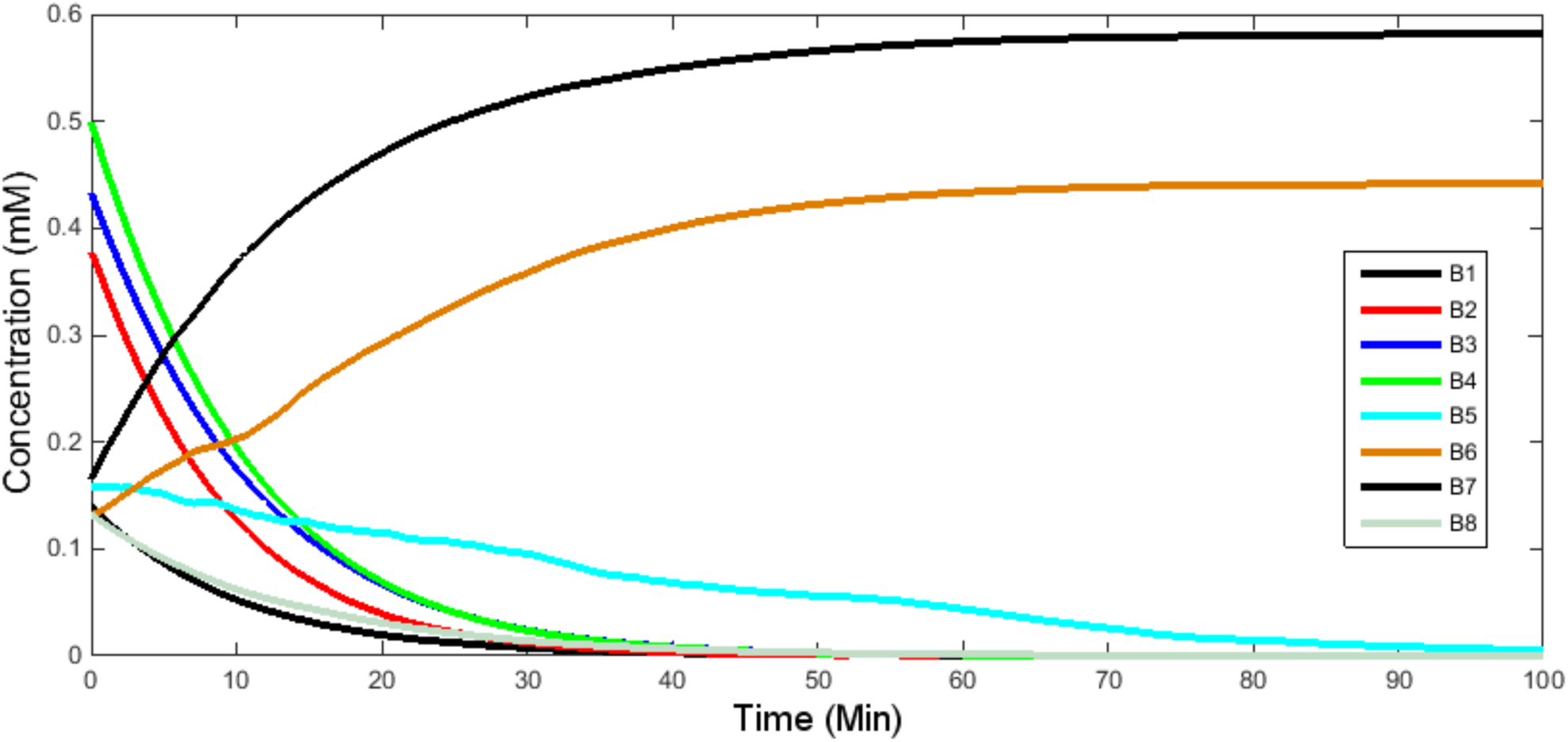
Concentration profiles for a number of both-direction metabolites. The results indicate that some of the metabolites are produced in the cells (i.e., metabolites with non-zero concentration) and some of the metabolites are consumed by the cells (i.e., metabolites with zero concentration).

### 2. Application to Glucose Blood Level

In the next simulation study, we evaluate the performance of the modelling framework in the presence of a disturbance. Generally speaking, the blood metabolite concentrations change due to a meal administration or an intravenous injection to the body. Here, in order to perform the robustness analysis, a single perturbation is introduced into the model during the simulation at a certain time. We have changed the blood glucose concentration I unit at time 40.

The simulation results illustrated in Figure 5 and Figure 6 show that once glucose concentration increases, the corresponding uptake flux increases as well (i.e., the absolute value of the glucose flux increases). In addition, the results indicate that glucose flux converges to zero once all glucose is depleted after a period of time (i.e., glucose is consumed by the cells).

**Figure 5:**
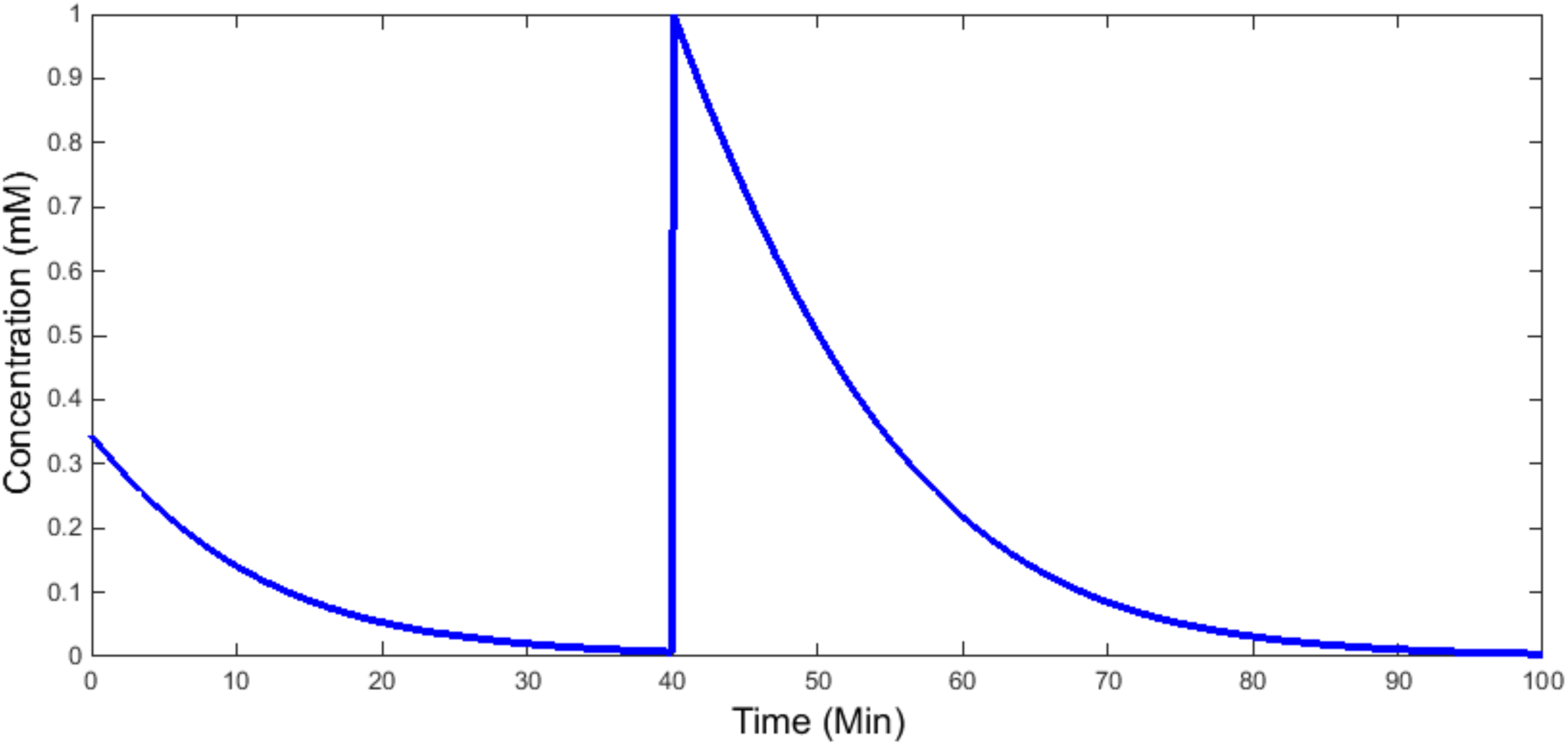
Concentration of glucose during a perturbation test. The result shows that the integration algorithm is robust with respect to perturbation in blood metabolites.

**Figure 6:**
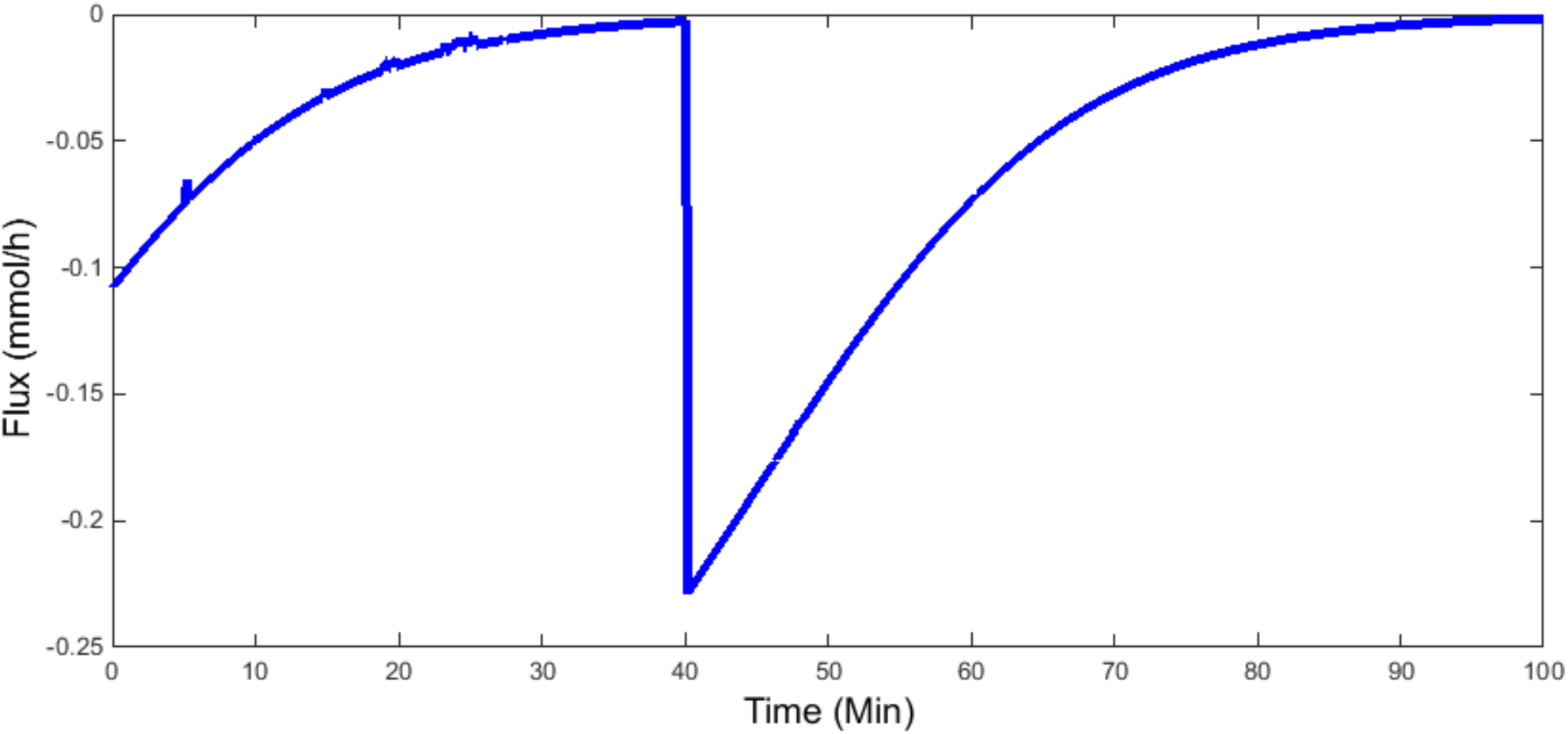
Corresponding glucose uptake rate during a perturbation test. The glucose uptake rate converges to zero because all glucose is consumed by the liver cell.

As seen, the developed framework is robust in the presence of perturbations in the blood metabolites. This opens up new opportunities to simulate multiple meal administrations and multiple intravenous injections, which has an important application in study design related to various diseases such as diabetes and obesity.

### 3. Application to Individual Metabolism Study

In the last simulation study, to demonstrate the impact of between individual variability in liver metabolism, we have designed an *in silico* study. In this study, we have created 50 subjects with different physiological properties (See Figure A.4). In addition, we have created 50 different sets of kinetic parameters associated with liver transport reactions indicating genetic variation between individual subjects (See Figures A.5 and A.6). In order to make a comparison for liver metabolism between individuals, we have simulated the model for certain time (i.e., 20 Minutes) and monitored the concentration of a consumed, produced and both-direction metabolite. In Figure 10, the simulation results are presented.

As seen in Figure 7, although simulations start with the same initial conditions for all subjects, there is significant difference in metabolite blood concentration levels after 20 minutes between subjects. This indicates the impact of genetic and physiological variability (i.e., Michaelis–Menten parameters and physiological properties variability) on the liver metabolism. In fact, this *in slico* study demonstrates the significance of between subject variability in human metabolism which is very important problem in personalized nutrition research community.

**Figure 7:**
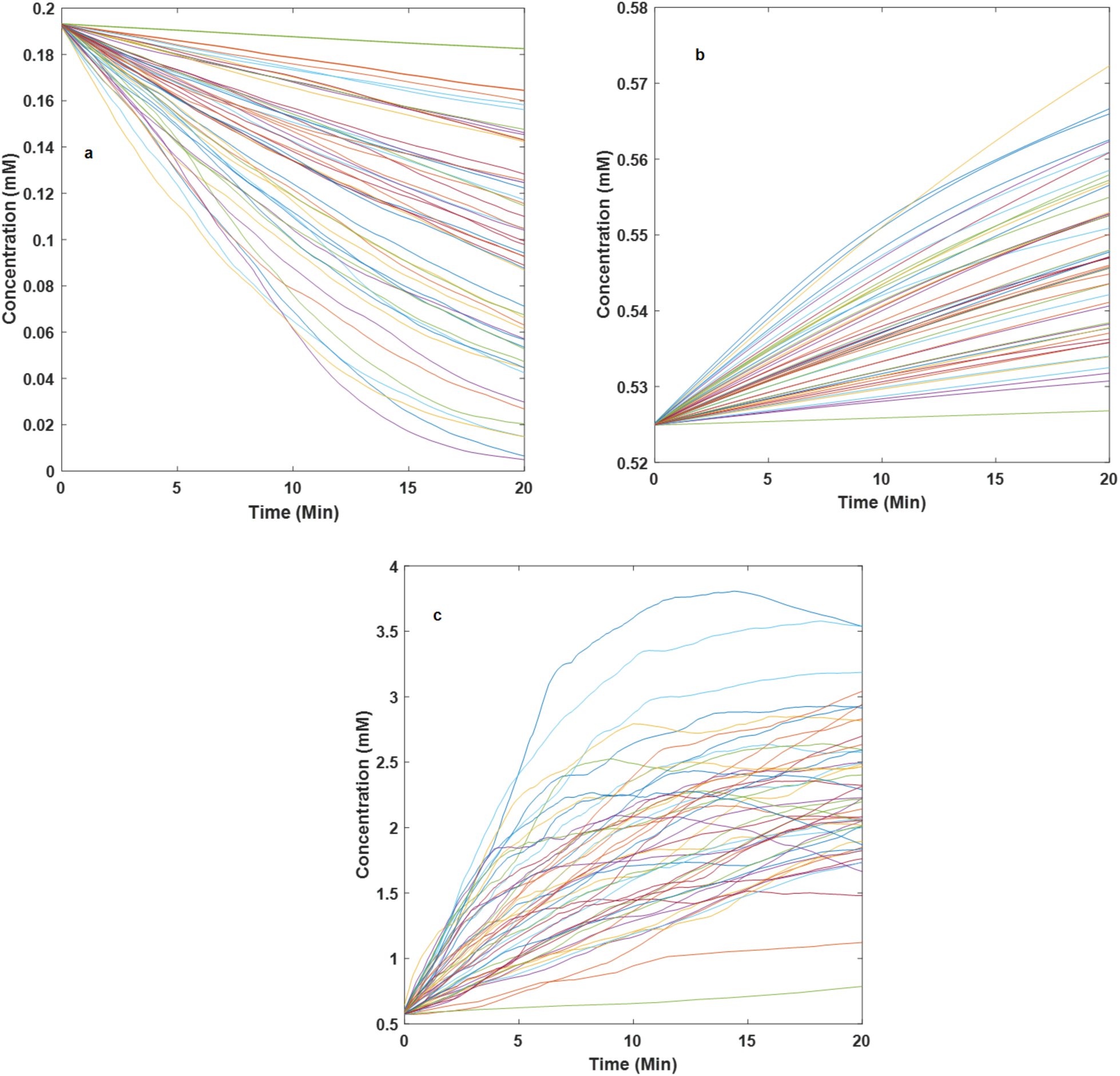
Time course of metabolite concentrations for 50 subjects. a) Consumed metabolite, b) produced metabolite, c) both-direction metabolite. The simulation results indicate that there is significant difference in blood metabolite concentration between individuals due to between individual variability (i.e., genetic and physiological differences).

## Conclusion

In this paper, we have developed a new modelling framework to capture multiple functions of the hepatocyte cells based on our previous multi-scale whole body metabolic model with a single objective function. The proposed framework has 3555 ODEs, including 1826 biochemical reactions. To evaluate the performance of the new model, simulations have been conducted with 74 objective functions, representing 74 exchange metabolites that can be produced by the liver cells. The simulation results indicate that the proposed modelling framework has promise with respect to convergence, computational efficiency and robustness. The developed model has several potential applications. For instance, it can be used to predict the blood metabolite concentrations after administrating multiple meals, to investigate the effect of multiple metabolic disorders on human metabolism and to investigate multiple hypotheses in personalized nutrition. However, there are some challenges. For example, several experimental studies should be carried out to have a reasonable approximation of substrate distribution between objective functions. In addition, the identification of the number of objective functions is a crucial step that requires a deep understanding of study designs and human cell physiology.

## Materials and Methods

To establish the modelling framework, our previous multi-scale whole body metabolic model has been used. In this model, a whole body model (WBM) with 15 compartments, including the lung, brain, muscle, gut, pancreas, liver, stomach, spleen, heart, bone, adrenal, skin, adipose tissue, and blood has been integrated with a human hepatocyte genome-scale model. The integration was performed using the dynamic flux balance analysis technique [17]. In addition, the whole body model contains 273 human serum metabolites. Therefore, the framework includes 3555 ordinary differential equations. The computational efficiency of the integration algorithm has been shown in our previous study [5]. Also, the applications of the modelling framework have been demonstrated to biomarker identification and blood alcohol concentration prediction in our previous studies [5, 6]. In both studies, a single objective function has been used in the parsimonious flux balance analysis problem to conduct the simulations. The mathematical representation of the multi-scale whole body metabolism with the dynamic flux balance analysis technique can be written as follows:

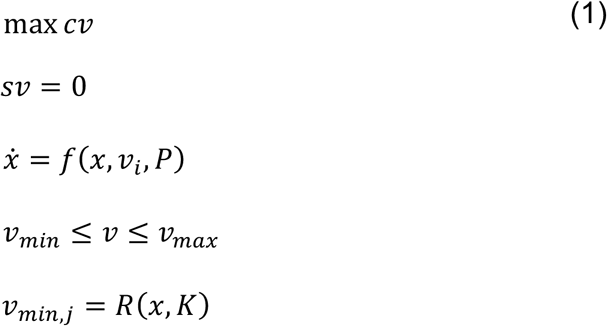

where *S* represents the stoichiometric matrix of the liver cell, *P* represents the model parameters, *R* represents the kinetic model, *v* represents reaction fluxes, *K* represents kinetic parameters, *x* represents metabolite concentrations and nonlinear function *f* represents the whole-body model.

In order to incorporate the multiple objective functions in the multi-scale whole body metabolism framework, we have included an additional constraint in the mathematical model (1) as follows:

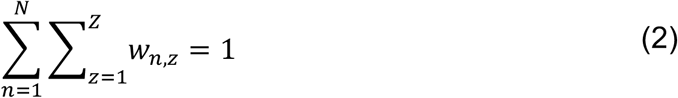

In Equation 2, *N* is the number of available substrates (i.e., consumed metabolites) and *Z* is the number of objective functions (for the specific model used here, Z=74). *w*_*n,z*_ is the fraction of substrate *n* that is used towards the objective function *z*. The interpretation of the added constraints is that the available substrates are divided based on the number of objective functions. Here, we assume that there is an objective function corresponding to every product that is synthesized by the liver cells (See Figure 8). For example, in Figure 8, we have two objective functions (i.e., P_1_ and P_2_) and one substrate (S). The consumed metabolite in this case is divided between each objective function with specific fractions, i.e., W_1_ and W_2_ such that W_1_+W_2_ becomes equal to W_T_. For this specific case, there were 163 metabolites that were consumed in the model.

**Figure 8:**
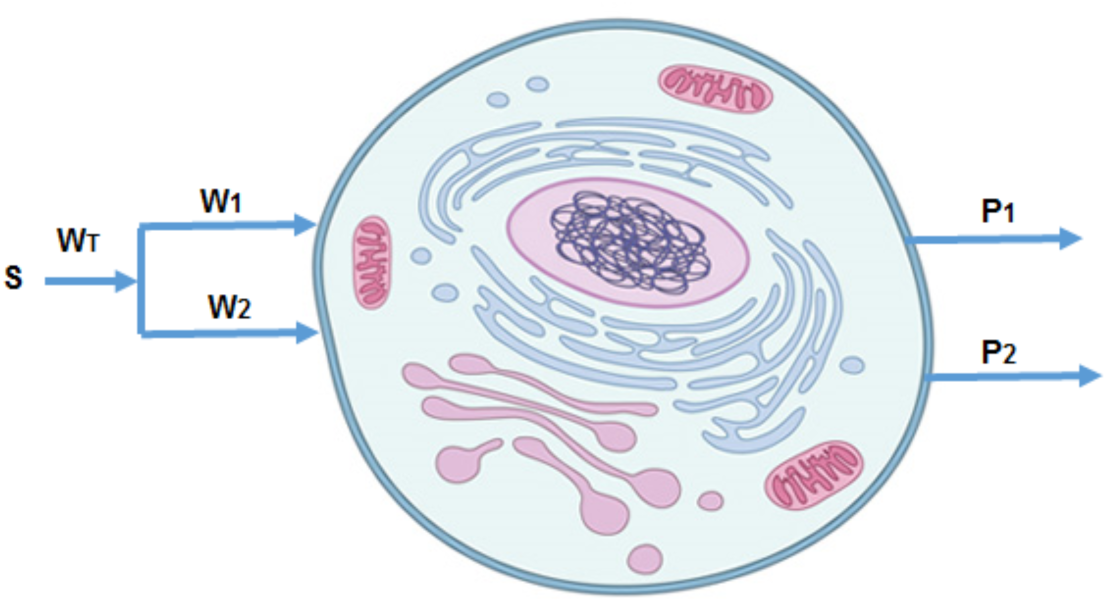
Interpretation of the constraint. In this example, the cell produces two metabolites (i.e., P_1_ and P_2_). The substrate (S) is available to make the products, and the substrate is divided between each objective function such that W_1_+W_2_ becomes equal to one. W_1_ is a fraction of the substrate (S) that is available for P_1_, and W_2_ is a fraction of the substrate that is available for P_2_, and W_1_+W_2_ equals to W_T_.

The mathematical formulation of the new multi-scale whole human body metabolism with multiple objective functions is written as follows:

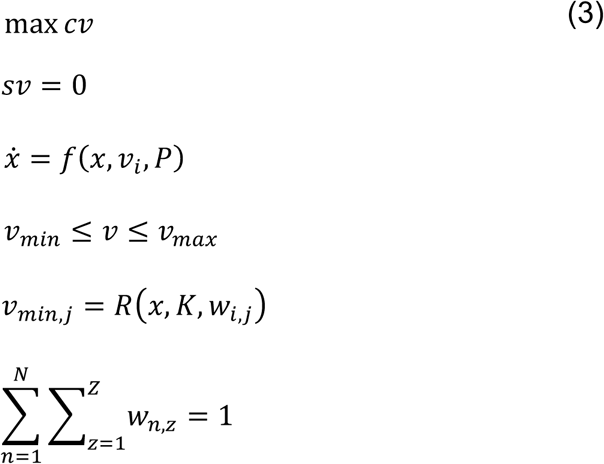

In fact, the fraction coefficients (i.e.,*w*_*n,z*_) can be chosen with a specific distribution or using experimental evidence. If a uniform distribution of these fractions are assumed, then the entire range of metabolic potential of the liver model can be comprehensively explored. Also, the number of objective functions can be selected such that all objectives cover main functions of the mammalian cells. Here, Michaelis Menten kinetics was assumed for all the 163 substrate uptake rates. The parameters were chosen in ad-hoc way (Figure A.5) and will be shown in the Supplementary Information upon publication.

In order to implement the developed modelling framework, dynamic flux balance analysis [17] is used. Following this algorithm, the initial conditions of the WBM are used to estimate the reaction rates. Then, the reaction rates are set as constraints for the FBA problem. The available substrates are divided into the number of objective functions in the FBA problem. Solving the FBA problem for each objective function leads to calculating metabolic fluxes throughout the entire network. Here, we assume the number of objective functions is equal to the number cell products (i.e., produced metabolites). The number of times that FBA is called in each time step is equal to the number of objective functions. Once the FBAs are solved for all objective functions, we need to calculate the net flux for each exchange reaction (i.e., the net flux is equal to the summation of all changes resulting from all FBA solutions). In the last step, the metabolite concentrations in the WBM are updated using net exchange fluxes. In this algorithm, we assume that the net fluxes are constant during each time interval and they are updated at the beginning of each time step. In Figure 9, the summary of the algorithm is presented. At the next section, we demonstrate the computational efficiency of implemented algorithm. All simulations are conducted using MATLAB and COBRA software.

**Figure 9:**
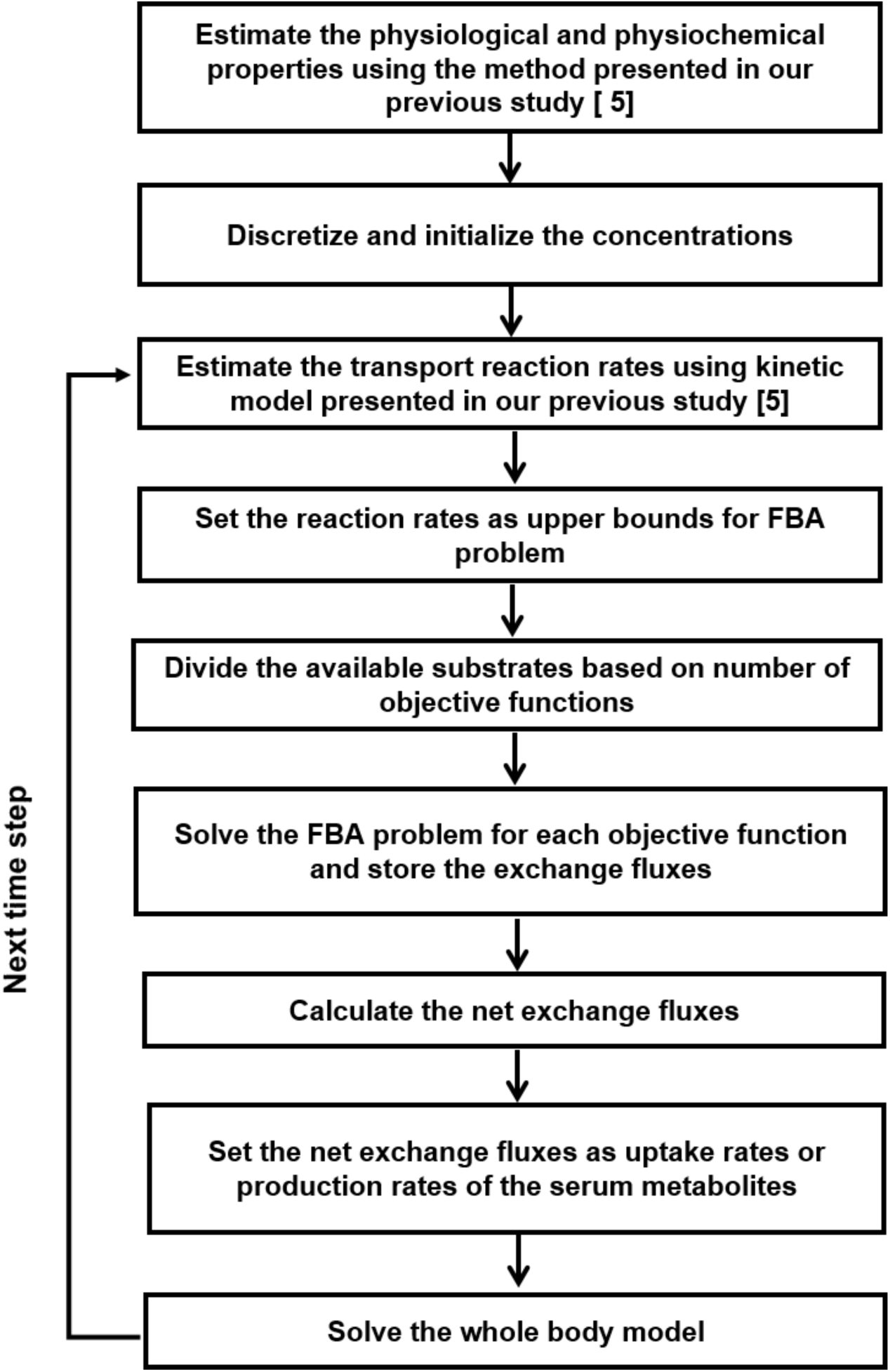
The algorithm for solving the integrated whole body metabolism with multiple objective functions. In this algorithm, we assume that the net fluxes are constant over each time step.

## Appendix

**Figure A.1:**
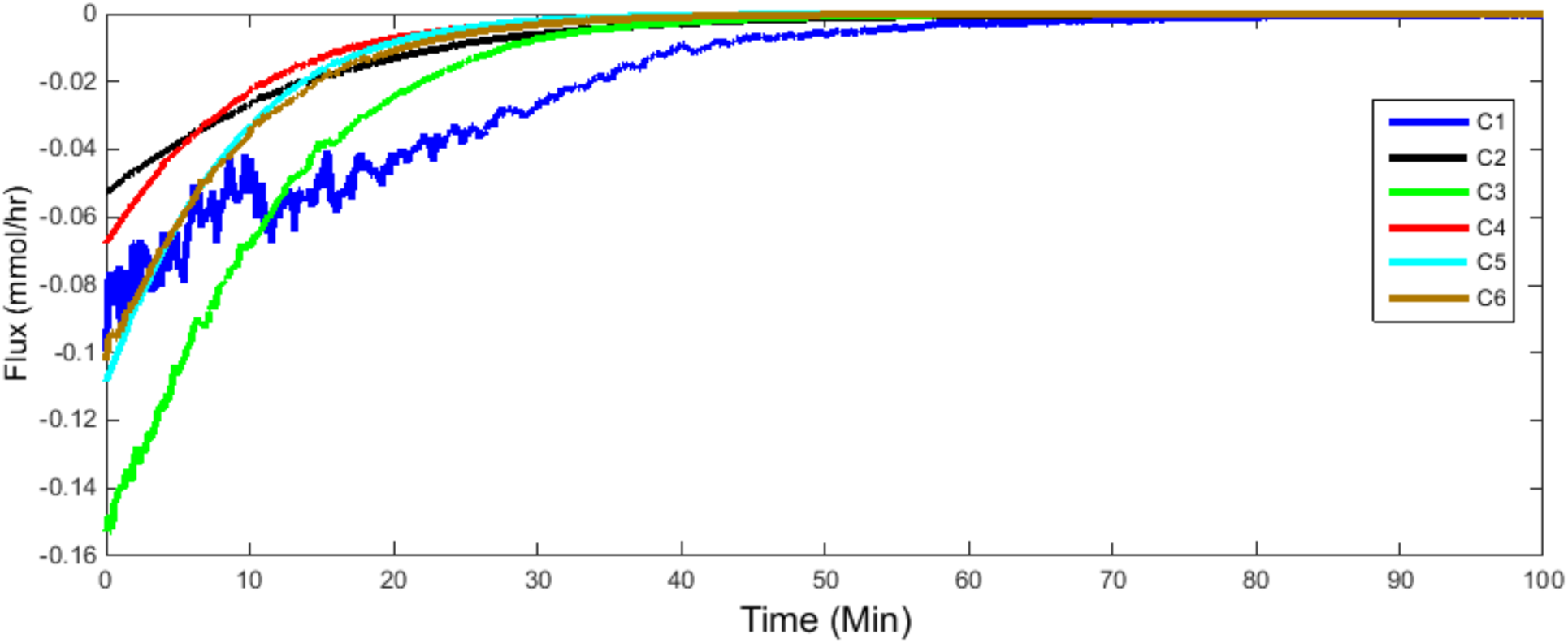
Corresponding flux profiles for a number of consumed metabolites. At steady-state conditions, the uptake fluxes approach zero.

**Figure A.2:**
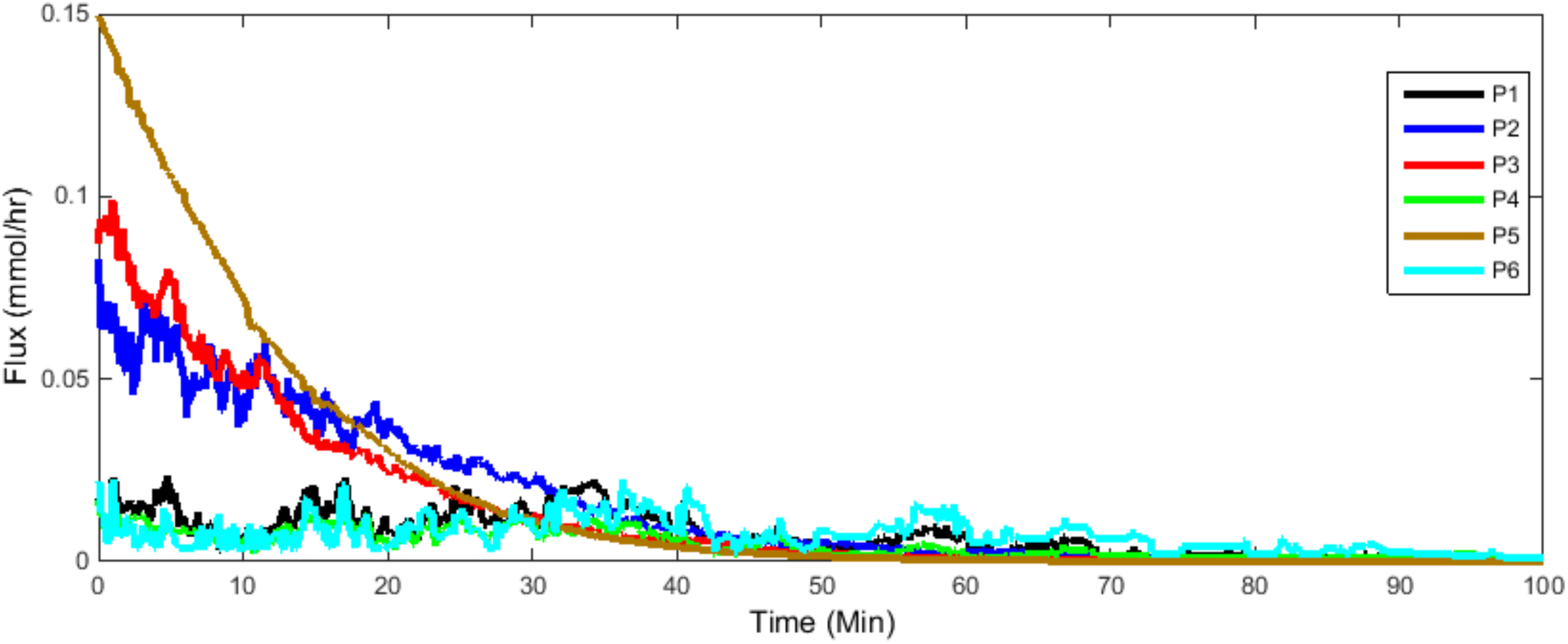
Corresponding flux profiles for a number of produced metabolites. At the steady-state conditions, the fluxes converge to zero.

**Figure A.3:**
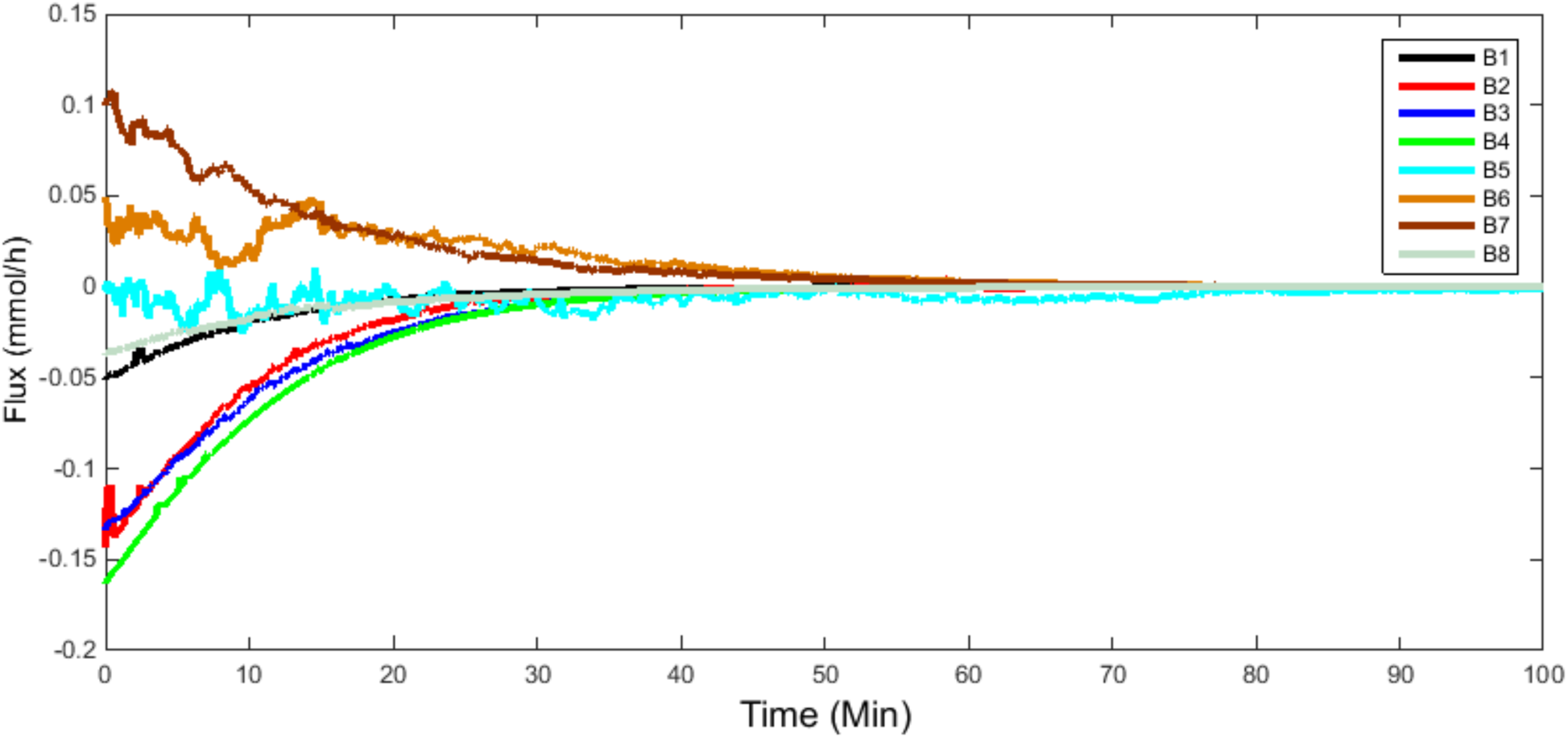
Corresponding flux profiles for a number of both-direction metabolites. The zero steady-state fluxes indicate the consistency between fluxes and concentrations.

**Table A.1:** Genome Scale Model of the Liver Cell

| **Property** | **Value** |
| --- | --- |
| Number of reactions | 1826 |
| Number of metabolites | 1360 |
| Number of exchange reactions | 237 |
| Total number of objective functions | 74 |

**Table A.2:**
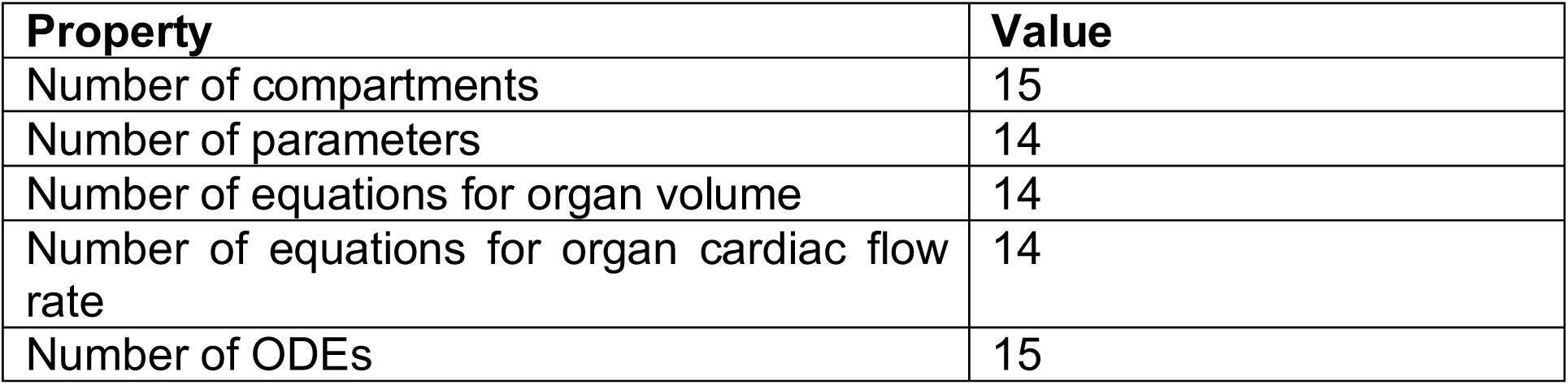
Whole-Human Body Model

**Table A.3:** Integrated Model

| **Property** | **Value** |
| --- | --- |
| Total number of compartments | 15 |
| Total number of exchange metabolites | 237 |
| Total number of parameters | 3318 |
| Total number of ODEs | 3555 |

**Figure A.4:**
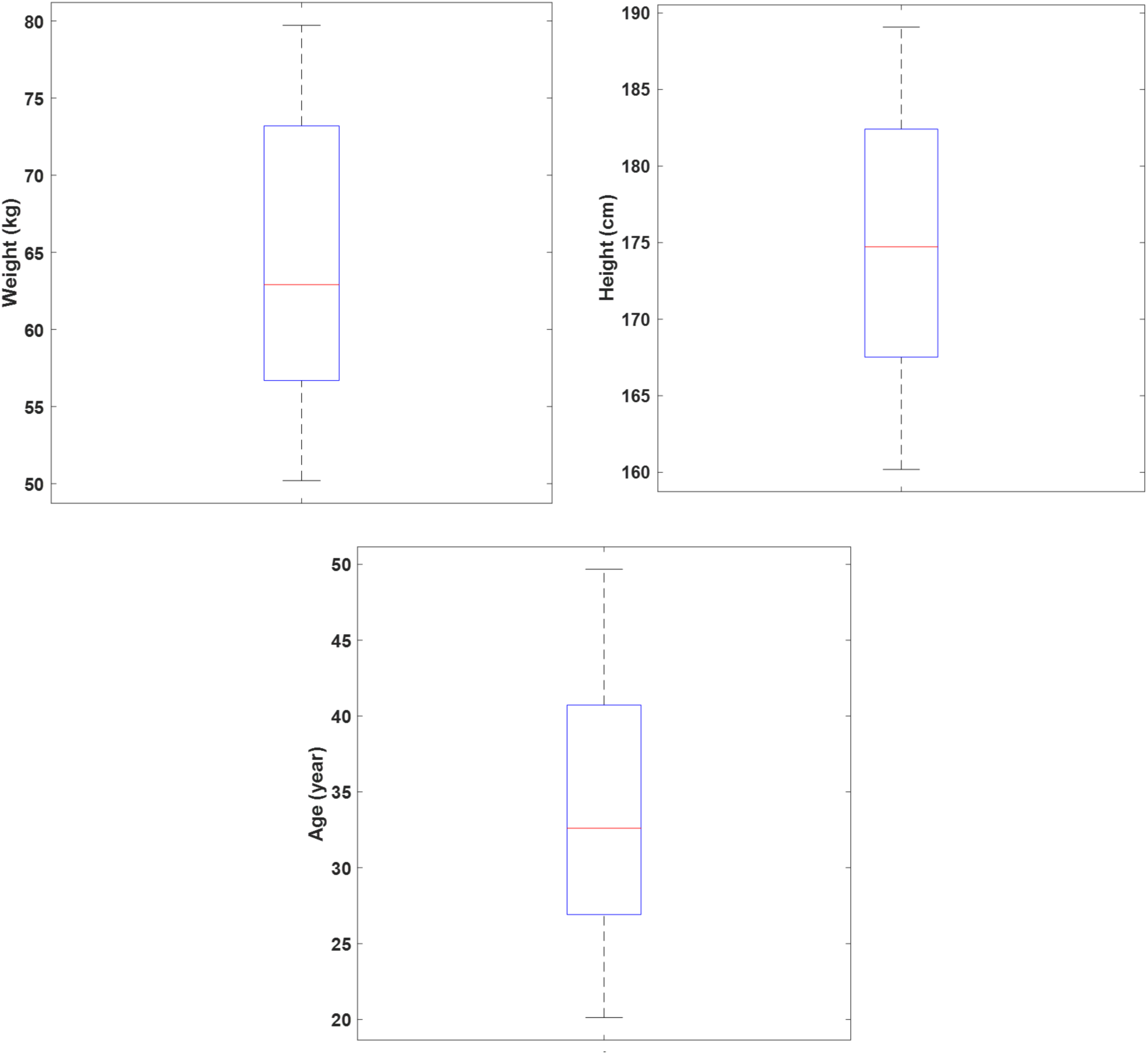
Physiological properties of the individual subjects

**Figure A.5:**
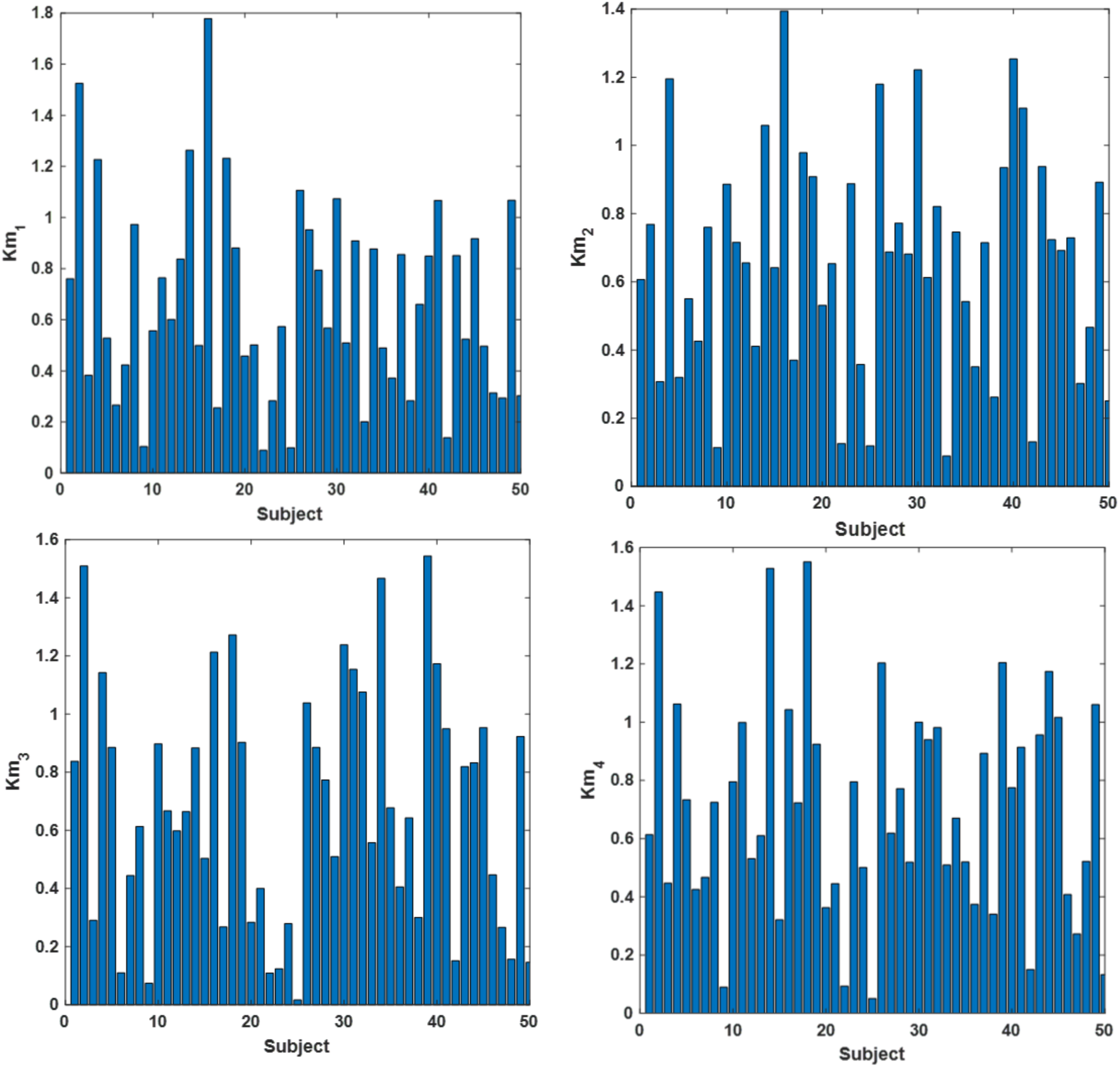
K_m_ Michaelis–Menten parameter for 50 subjects. Here, the bar charts show the values for only 4 transport reactions, however there are 163 km values since the model includes 163 consumed metabolites.

**Figure A.6:**
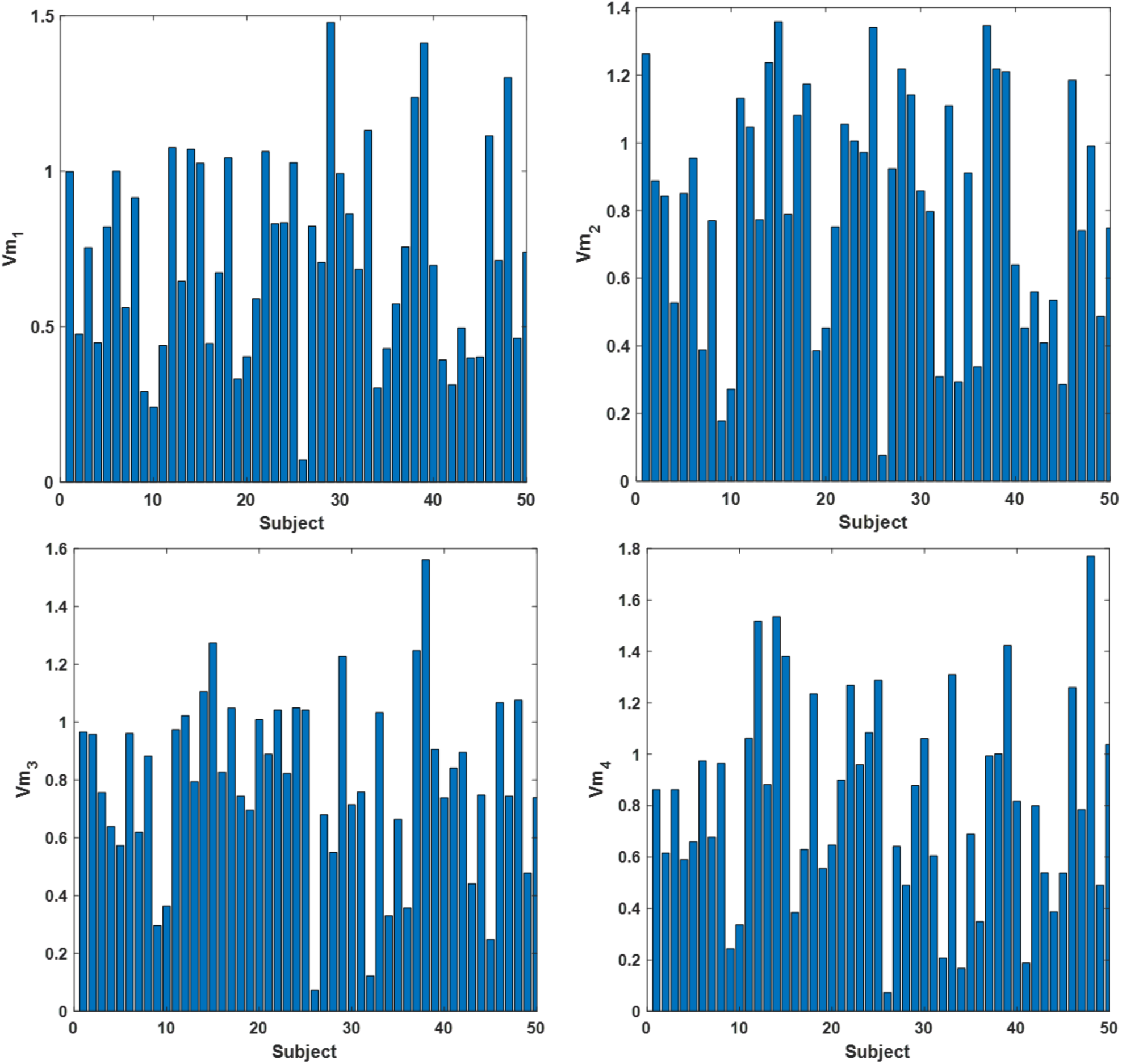
V_m_ Michaelis–Menten parameter for 50 subjects. Here, the bar charts show the values for only 4 transport reactions, however there are 163 V_m_ values since the model includes 163 consumed metabolites.

